# Near real-time data on the human neutralizing antibody landscape to influenza virus as of early 2026 to inform vaccine-strain selection

**DOI:** 10.64898/2026.02.18.706711

**Authors:** Caroline Kikawa, John Huddleston, Sam A. Turner, Andrea N. Loes, Jiaojiao Liu, Sydney Gang, Tachianna Griffiths, Elizabeth M. Drapeau, Benjamin J. Cowling, Faith Ho, Nancy H. L. Leung, Janet A. Englund, Kirsten Lacombe, Shinji Watanabe, Hideki Hasegawa, Michael Busch, Marion Lanteri, Mars Stone, Bryan Spencer, Richard A. Neher, Derek J. Smith, Trevor Bedford, Scott E. Hensley, Jesse D. Bloom

**Affiliations:** Division of Basic Sciences and Computational Biology Program, Fred Hutch Cancer Center, Seattle, WA; Department of Genome Sciences, University of Washington, Seattle, WA; Medical Scientist Training Program, University of Washington, Seattle, WA; Vaccine and Infectious Disease Division, Fred Hutch Cancer Center, Seattle, WA; Howard Hughes Medical Institute, Seattle, WA; Center for Pathogen Evolution, Department of Zoology, University of Cambridge, Cambridge, UK; Depts of Microbiology and Medicine, Perelman School of Medicine, University of Pennsylvania, Philadelphia, PA; WHO Collaborating Centre for Infectious Disease Epidemiology and Control, School of Public Health, LKS Faculty of Medicine, The University of Hong Kong, Pok Fu Lam, Hong Kong Special Administrative Region; Seattle Children’s Research Institute and Department of Pediatrics, University of Washington, Seattle, WA; Influenza Research Center, National Institute of Infectious Diseases, Japan Institute for Health Security, 1-21-1 Toyama Shinjuku-ku, Tokyo 162-8655, Japan; Vitalant Research Institute, San Francisco, CA; Department of Laboratory Medicine, University of California San Francisco, San Francisco, CA; Creative Testing Solutions, Tempe AZ; American Red Cross, Dedham, MA; Swiss Institute of Bioinformatics and Biozentrum, University of Basel, Basel, Switzerland

## Abstract

Twice each year, a decision is made on whether to update the strains included in the seasonal influenza vaccine to better match the most recent circulating viral strains. To characterize the antigenic properties of current seasonal influenza A strains to inform the upcoming decision about which strains to include in the 2026-2027 Northern Hemisphere vaccine, here we perform high-throughput sequencing-based neutralization assays using a library of 57 H3N2 and 34 H1N1 influenza hemagglutinins reflecting the circulating diversity of strains in late 2025 to early 2026. We assay this library against 302 human sera collected in late 2025. The resulting data set encompasses 27,409 titers, and provides a near real-time portrait of the human neutralizing antibody landscape against influenza virus. We find that many human sera have lower titers against the K subclade of H3N2 and the D.3.1.1 subclade of H1N1; these subclades have recently become dominant among their respective subtypes. Our measurements also reveal variability in titers to different subvariants within the K subclade of H3N2, with titers especially low to subclade K strains with additional mutations in antigenic regions D and E. We make all our data and accompanying visualizations publicly available to enable their use in vaccine-strain selection and analyses of influenza evolution and immunity.

## Introduction

The hemagglutinin (HA) proteins of seasonal influenza viruses continuously evolve to erode human neutralizing antibody immunity such that people are re-infected with an influenza A virus roughly every 5 years (Ranjeva et al. 2019; Kucharski et al. 2018). The influenza A subtypes that co-circulate in humans, H3N2 and H1N1 (descended from the 2009 H1N1 pandemic strain), acquire 3-4 and 2-3 amino acid substitutions in their HA proteins per year, respectively (Bedford et al. 2014; Smith et al. 2004).

To keep pace with this evolution, updates to the strains in the seasonal influenza vaccine are considered biannually with the goal of ensuring the vaccine strains are well matched to circulating strains. Recommendations of which strains to include in updated vaccines are made at vaccine-composition meetings, with a meeting in late winter to choose the strain for the upcoming Northern Hemisphere influenza season, and a meeting in early autumn to choose the strain for the upcoming Southern Hemisphere influenza season (World Health Organization 2024, n.d.-a, n.d.-b). The recommendations are based on a combination of viral sequence data reflecting the relative prevalence of different viral strains in the human population, and serological data quantifying the antigenic properties of strains (Luksza and Lässig 2014; Neher et al. 2016; Huddleston et al. 2020; Shi et al. 2025). Historically, the serological data consisted of hemagglutination inhibition assays performed with sera from ferrets infected with defined viral strains (Smith et al. 2004; Jorquera et al. 2019). However, there has been increasing use of measurements made using human sera (World Health Organization 2024, n.d.-a, n.d.-b; Ampofo et al. 2015; Fonville et al. 2016; Kikawa et al. 2025) as there is growing recognition that the neutralizing antibody specificities of humans are more complex than that of singly infected ferrets (Fonville et al. 2016; Linderman et al. 2014; Cobey and Hensley 2017; J. M. Lee et al. 2019a).

Following a period of low influenza genetic diversity during and after the COVID-19 pandemic, influenza evolution prior to the September 2025 vaccine-strain selection meeting was characterized by the rapid emergence and spread of multiple H3N2 variants carrying mutations at important antigenic sites (Sabaiduc et al. 2025; Huddleston et al. 2025). The September 2025 meeting recommended for the 2026 Southern Hemisphere vaccine an updated H3N2 strain from subclade J.2.4 and an updated H1N1 strain from subclade D.3.1 (World Health Organization, n.d.-b) (throughout this paper we use a new dynamic nomenclature system for influenza subclades (Neher et al. 2026)). However, by the end of the 2025, additional antigenically divergent daughter subclades had emerged and become predominant for both subtypes: H3N2 subclade K (derived from J.2.4 with HA1 mutations K2N, S144N, N158D, I160K, Q173R, and T328A) and H1N1 subclade D.3.1.1 (derived from D.3.1 with HA1 mutations R113K, A139D, E283K, and K302E). Very recent experimental studies have shown that subclade K is antigenically distinct (Liu et al. 2026; Wang et al. 2026; Dee et al. 2026; Kirsebom et al. 2025; Separovic et al. 2026; Ikonen et al. 2026; Cheng et al. 2026), with human sera having ∼2-fold lower titers to subclade K than other recent H3N2 strains (Liu et al. 2026; Wang et al. 2026), although the exact magnitude of the titer decrease varied across cohorts and studies (Wilson et al. 2026; Guiomar et al. 2026). Additional HA mutations have since arisen in subvariants of both subclades K and D.3.1.1, although the effects of these new mutations have not been experimentally characterized prior to the current study.

Here we characterize the neutralization by human sera of current human seasonal H3N2 and H1N1 strains to provide data to inform the upcoming decision of which strains to include in the 2026-2027 Northern Hemisphere vaccine. We do this by using a recently developed sequencing-based neutralization assay (Kikawa et al. 2025; Loes et al. 2024; Kikawa et al. 2026) that can rapidly measure titers of many human sera against many viral strains. We previously used this assay to generate data prior to the September 2025 vaccine-strain selection decision (Kikawa et al. 2025). This study takes a similar approach but uses an updated set of H3N2 and H1N1 strains representative of those currently circulating in the human population (including multiple subvariants with subclades K and D.3.1.1) and a new set of human sera collected mostly in October to November 2025. We measure a total of 27,409 neutralization titers to provide a near real-time portrait of human neutralizing antibody immunity to seasonal influenza.

## Results

### A library of influenza HAs representing the diversity of human H3N2 and H1N1 influenza in late 2025 to early 2026

Our goal was to design a library of influenza HAs that could be used to measure neutralization titers of human sera against recently circulating influenza HAs prior to the February 2026 vaccine-strain selection (**Figure 1**). In November 2025, we examined all available HA sequences from human H3N2 and H1N1 strains, and chose a set of naturally occurring human seasonal HAs with the goal of covering the existing HA diversity. Specifically, we chose HAs from recently circulating strains with high frequency over the last six months window, strains that included mutations at previously-defined antigenic or receptor binding site-adjacent sites (Koel et al. 2013; Wolf et al. 2006; Caton et al. 1982), strains that appeared to be rapidly increasing in frequency (Abousamra et al. 2024), and strains with mutations that had arisen more recurrently than expected (Bloom and Neher 2023).

**Figure 1.**
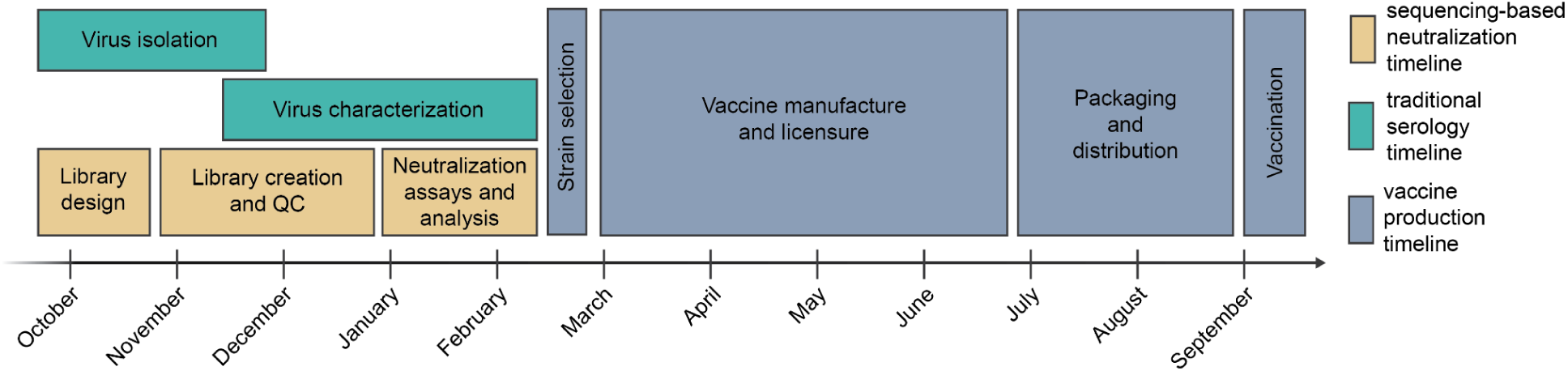
Typical timeline for vaccine-strain selection and production for the Northern Hemisphere influenza season. A decision about which strains to include in the annual vaccine is made twice each year: typically in early autumn for the Southern Hemisphere vaccine that will be used in the upcoming summer, and in late winter for the Northern Hemisphere vaccine that will be used in the upcoming autumn and winter. Illustrated above is the typical timeline for influenza vaccine strain selection and production for the Northern Hemisphere influenza seasons. The periods for traditional virus isolation and characterization are shown alongside our timeline for the sequencing-based neutralization assay measurements reported in the current study.

Overall our library included 53 recently circulating human H3N2 strains and 30 recently circulating human H1N1 strains (**Figure 2** and **Supplementary File 1**). As of early 2026, the HAs we chose continue to cover most of the diversity of the HAs of sequenced human H3N2 and H1N1 influenza (**Figure 2**). Additionally, we included HAs from eight vaccine strains dating back to the 2020 vaccine for H3N2 and the 2018 vaccine for H1N1.

**Figure 2.**
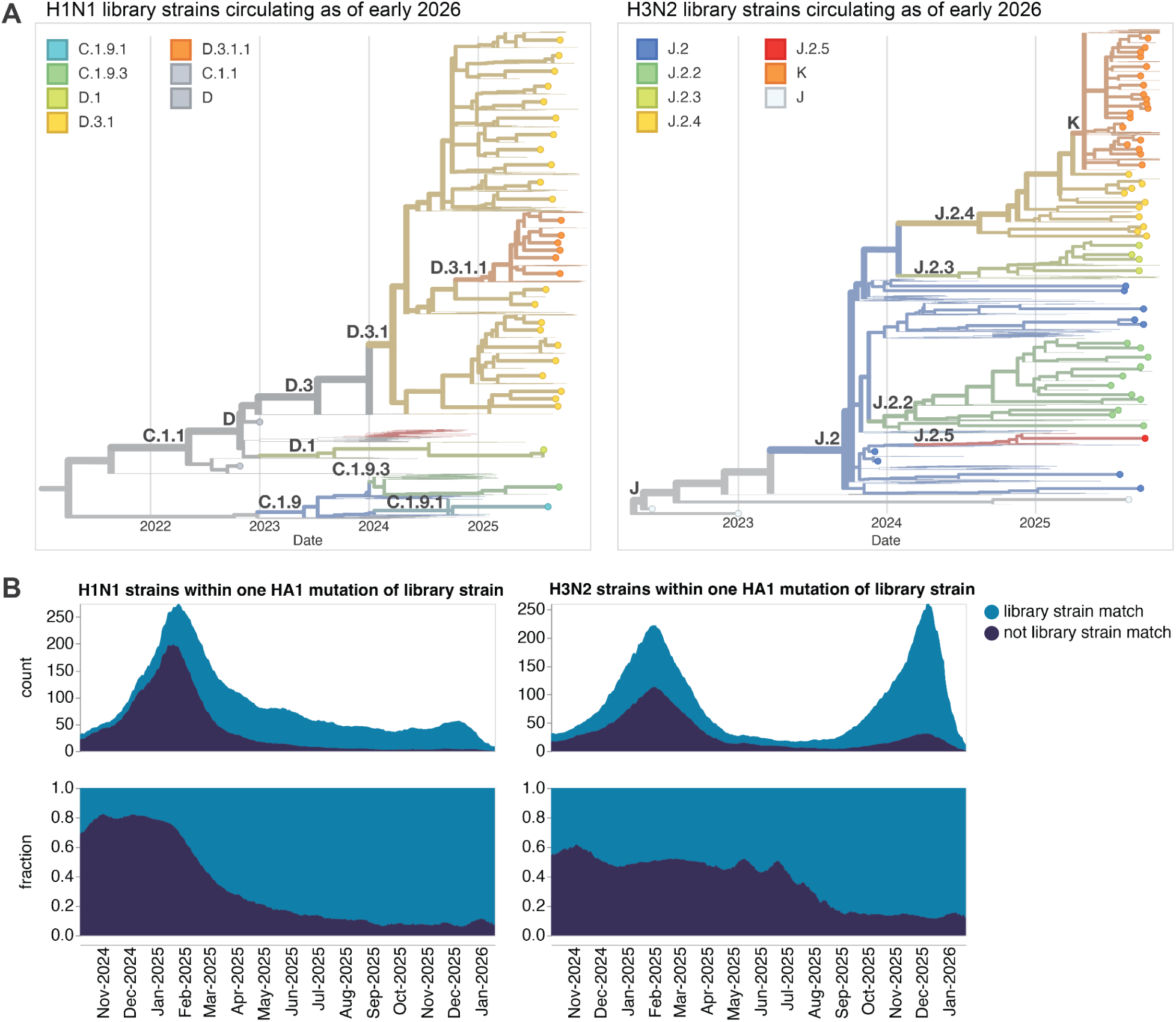
The HA sequences chosen for our sequencing-based neutralization library are representative of human seasonal influenza circulating in late 2025 and early 2026. (**A**) Phylogenetic trees of HA genes of H1N1 and H3N2 strains chosen for our sequencing-based neutralization assay library are shown as points, with other recently circulating strains shown as thin lines. Strains are colored by their subclade designation. Interactive versions of these trees are at https://nextstrain.org/groups/blab/kikawa-seqneut-2025-2026-VCM/h1n1pdm and https://nextstrain.org/groups/blab/kikawa-seqneut-2025-2026-VCM/h3n2. (**B**) Count and fraction of all human seasonal H1N1 and H3N2 HA sequences available as of 2026-02-05 that closely match a strain in our sequencing-based neutralization assay library (within one HA1 amino-acid mutation). Counts and fractions are averaged over a sliding 10-day window.

We next generated barcoded viruses expressing each of the chosen HAs with the other viral genes from the lab-adapted A/WSN/1933 strain, using previously described approaches (Loes et al. 2024; Kikawa et al. 2026). Most HAs were associated with two or three distinct barcodes to provide internal replicates in the sequencing-based neutralization assays. After quality control, we had a total of 186 unique barcoded viral variants for the 91 different HAs.

### A panel of sera collected in late 2025 from humans from multiple locations

We assembled a collection of 302 sera from individuals ranging from 0 to 103 years of age from five different locations (see **Figure 3**, which defines the abbreviations we use to refer to each sera set). Most sera were collected in October 2025-November 2025, with a few sera from the HKU set collected earlier in 2025. The PENN sera were collected pre- and post-vaccination with the Northern Hemisphere 2025-2026 seasonal influenza vaccine (Flulaval Trivalent), which contained a J.2 clade H3N2 component and a D clade H1N1 component; note that some of these sera have recently been analyzed against a few H3N2 viral strains by hemagglutination-inhibition assays (Liu et al. 2026). We included the post-vaccination sera to quantify how the current vaccine affects titers against current circulating strains; however, it is thought that at a global population level, antibody titers are shaped more by infection than vaccination (Davis et al. 2020; J. S. Turner et al. 2020). Some of the HKU and NIID sera are also from individuals with information on recent infection or vaccination status (**Supplementary File 2**). Most of the remaining sera are from individuals with unknown or incomplete infection and vaccination histories. We prioritized sampling sera from many unique individuals from wide age ranges and geographical locations because these factors are thought to contribute to person-to-person differences in neutralizing antibody titers (Cobey and Hensley 2017).

**Figure 3.**
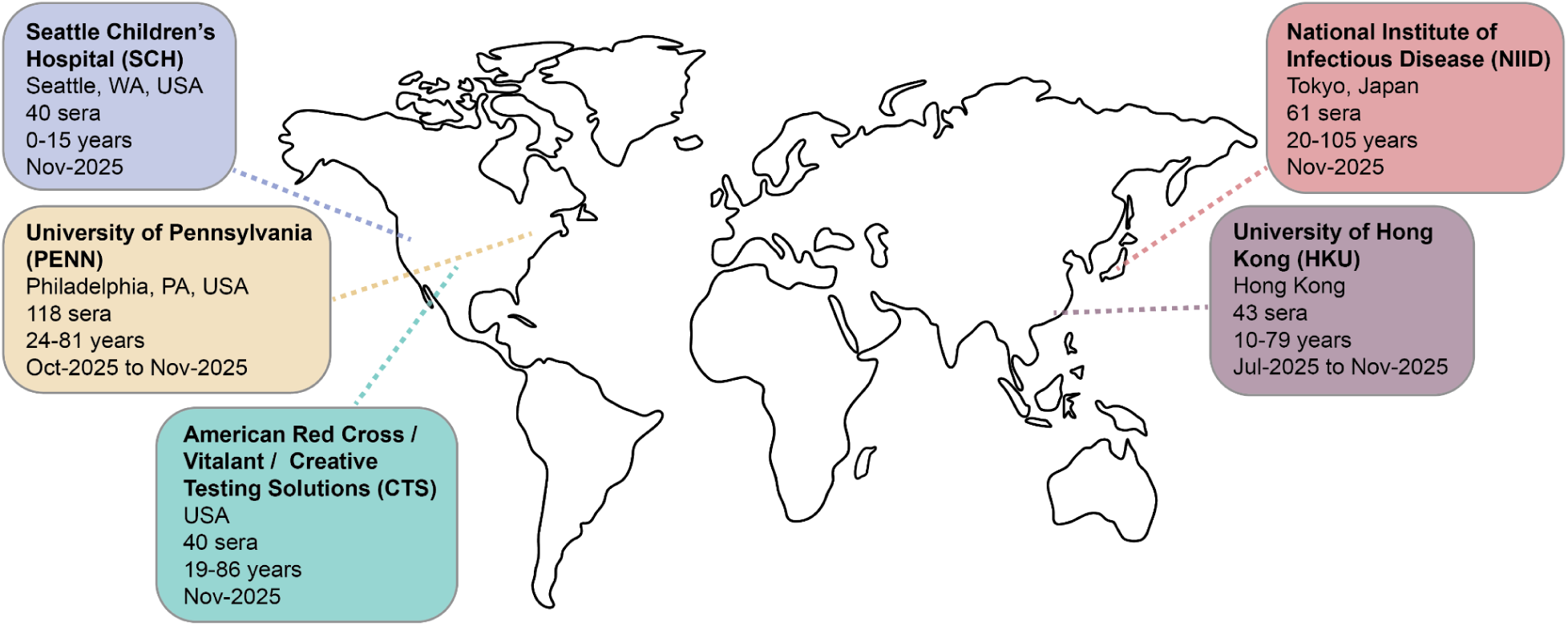
Overview of the human sera used in the neutralization assays. The map above summarizes the general location, number, distribution of ages, and timeframe of collection for the five sets of human sera used in this study. All the sera were taken from unique individuals with the exception of the University of Pennsylvania sera, for which we tested matched pre- and post-vaccination sera from 59 unique individuals.

### Neutralizing antibody landscapes to recent H3N2 influenza strains

We performed sequencing-based neutralization assays to measure neutralization of the 91 viral strains against all 302 sera, for a total of 27,409 titers (a small fraction of titers were dropped due to low-quality neutralization curves; see **Methods**). We quantify the titers as the reciprocal serum dilution that neutralizes 50% of the infectivity of a given viral strain. In this section we describe the 17,142 titers measured against the H3N2 strains.

The median titers across all sera differed by more than three-fold across the recent human H3N2 strains (**Figure 4**, **Figure 5**, and interactive plots linked in those figure legends). The strains with the lowest titers were in subclade K, a finding consistent with recent studies (Liu et al. 2026; Wang et al. 2026; Dee et al. 2026; Kirsebom et al. 2025; Separovic et al. 2026) showing that this rapidly growing subclade (which is now dominant among H3N2) is neutralized less well by human sera than the both the J.2:S145N strain in the current 2025-2026 Northern Hemisphere vaccine and the J.2.4 strain chosen in September 2025 for the 2026 Southern Hemisphere vaccine (**Figure 4**). Across subclades, titers to subclade K strains were significantly lower than all other non-subclade K strains in the library; when compared to each other subclade individually, titers to subclade K strains were not significantly different from J.2.4 strains but were significantly lower than J.2, J.2.2 and J.2.3 strains (**Supplementary Figure 1A,B**).

**Figure 4.**
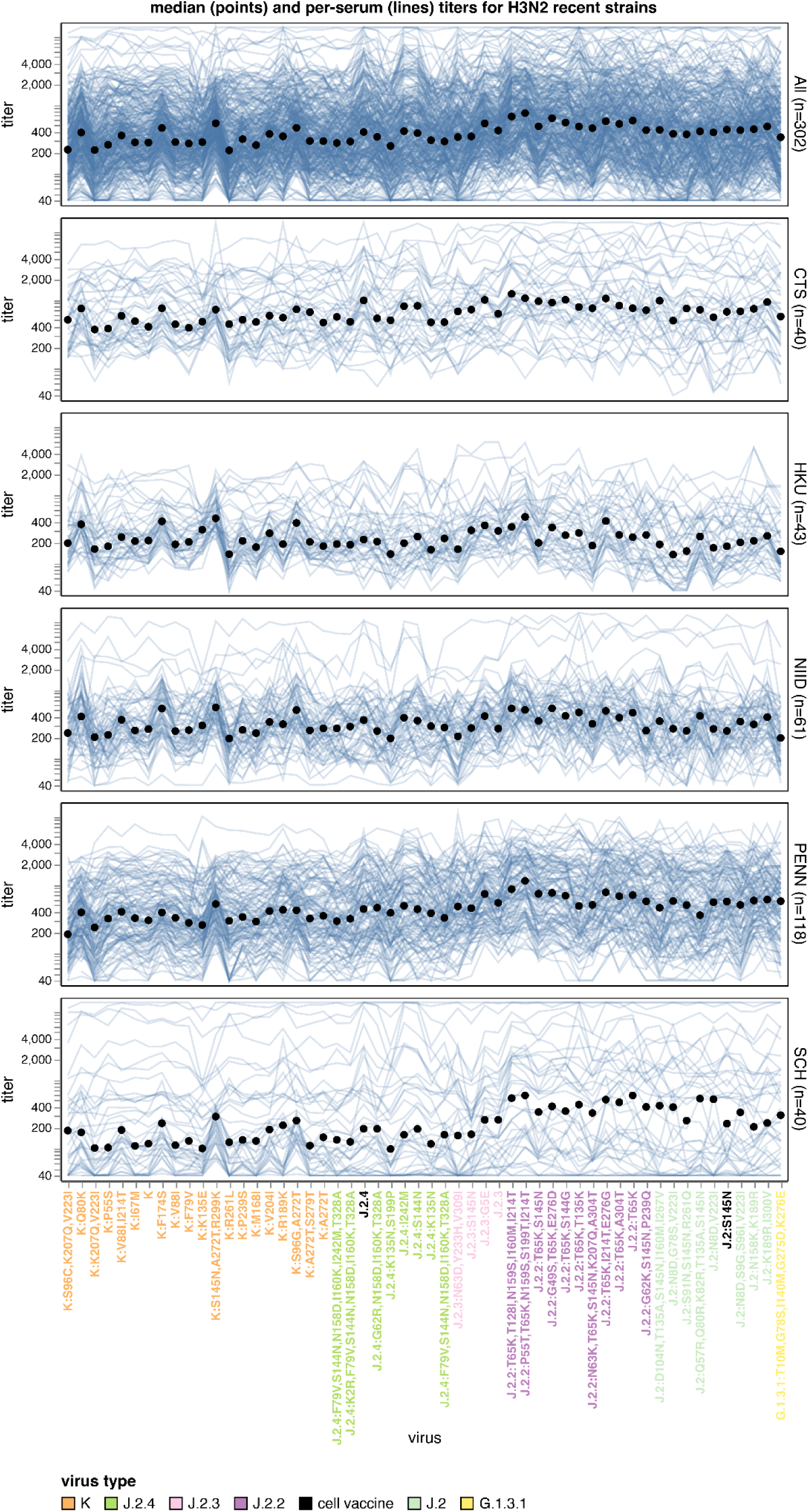
Human neutralizing antibody landscape against recent H3N2 viruses. Median titer across sera (black points) and titers for individual sera (blue lines) against the recently circulating H3N2 strains in the library. The top plot shows titers for all 302 sera, and the other plots show titers by serum set. The viral HA1 haplotype labels are colored by their subclade. The J.2:S145N strain is the current cell-based 2025-2026 Northern Hemisphere vaccine strain, and a J.2.4 strain was chosen in September 2025 for the cell-based 2026 Southern Hemisphere vaccine (these strains are labeled in black). The dynamic range of our assays (determined by the dilution series used for the sera) enabled measurements of titers between 40 and 13619; sera with titers above or below this range are censored accordingly. See https://jbloomlab.github.io/flu-seqneut-2025to2026/human_H3N2_recent_individual_sera_vertical.html for an interactive version of this plot that can be subset on specific sera or age groups, and for which you can mouseover points and lines for details about viruses and sera. See https://jbloomlab.github.io/flu-seqneut-2025to2026/human_H3N2_recent_interquartile_range_vertical.html for a comparable plot showing the median and interquartile range. Note that this figure shows just the titers against the recent strains and the 2025-2026 and 2026 vaccine strains (54 strains total); see https://jbloomlab.github.io/flu-seqneut-2025to2026/ for plots that show titers against the older vaccine strains.

**Figure 5.**
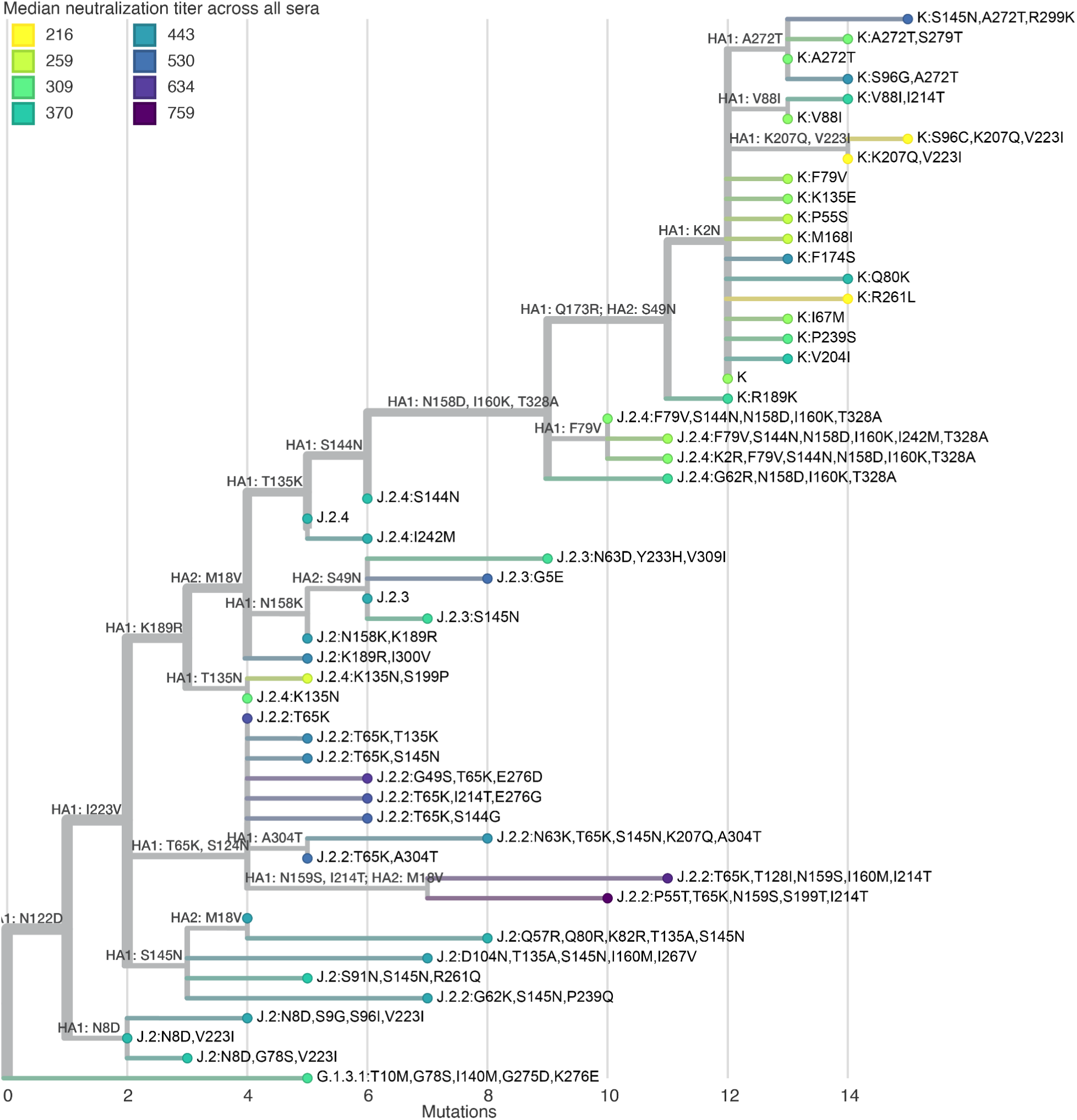
Phylogenetic tree of H3N2 HA proteins colored by median titer across all sera. Tree of H3N2 HA sequences in the library colored by the median titer across all sera. The tree is built on the protein sequences and branch lengths are amino-acid mutations relative to the root. All HA1 and HA2 mutations are labeled on branches. See https://nextstrain.org/community/jbloomlab/flu-seqneut-2025to2026@main/H3N2 for an interactive version of this tree with additional coloring options, a measurements panel with the individual serum titers, and an option to show all amino-acid mutations on branches.

All sera sets also show appreciable variation in titers among different subclade K strains. The subclade K strains with the lowest titers have mutations in the defined antigenic regions (Wu and Wilson 2017; M.-S. Lee and Chen 2004) D (sites 96, 207, and 223) or E (site 261) (**Figure 4**, **Figure 5**, and **Supplementary Figure 1C**). This observation is important because the canonical subclade K strain has many mutations in antigenic regions A and B but fewer mutations in regions D and E (Liu et al. 2026); our results suggest that neutralization of subclade K is further eroded by mutations in regions D and E, suggesting strains with such mutations could spread in the future. There are also a handful of strains from J.2.4 with mutations to 135N (which adds a potential N-linked glycosylation site) that have titers comparably low to some subclade K strains (**Figure 4**, **Figure 5**, and **Supplementary Figure 1D**).

In addition to the aforementioned trends in the median titers across sera, there is dramatic variation across different sera (**Figure 4**, especially see interactive version linked in figure legend, which makes it easier to explore individual sera). Some of this variation just represents sera that have higher or lower overall titers to all strains. But inspection of the lines in the interactive version of **Figure 4** identifies sera that are strongly impacted by specific mutations: for instance, some sera have dramatically altered titers to subclade K strains with K135E or R189K mutations (both of which are antigenic sites that have undergone recent evolution (Sabaiduc et al. 2025)), although these trends are not apparent in aggregated titers since the mutations affect only a subset of sera (**Supplementary Figure 1C**). We can also identify some trends that stratify by age group. For instance, sera from children (the SCH set from Seattle Children’s Hospital) have a much more pronounced trend than sera from adults (the other sera sets) towards lower titers to the more recent J.2.3, J.2.4, and K subclades versus the J.2, J.2.2, and G.1.3.1 subclades (**Figure 4**).

### Neutralizing antibody landscapes to recent H1N1 influenza strains

The 10,267 titers that we measured for the 302 human sera against the H1N1 strains are shown in **Figure 6A** and **Figure 7**. The variation in median titers among H1N1 strains is about two-fold, which is somewhat lower than that among the H3N2 strains. Nearly all the H1N1 strains with the lowest median titers are in subclade D.3.1.1 (daughter clade to D.3.1 with HA1 mutations R113K, A139D, E283K and K302E), which has become dominant among H1N1 strains over the last six months. Titers to all six subclade D.3.1.1 strains in our library are lower than the titers to both the subclade C.1.1 strain in the current 2025-2026 Northern Hemisphere vaccine and subclade D.3.1 strains, including the strain chosen in September 2025 for the 2026 Southern Hemisphere vaccine strain (**Figure 6A**, **Figure 7**, and **Supplementary Figure 1E).**

**Figure 6.**
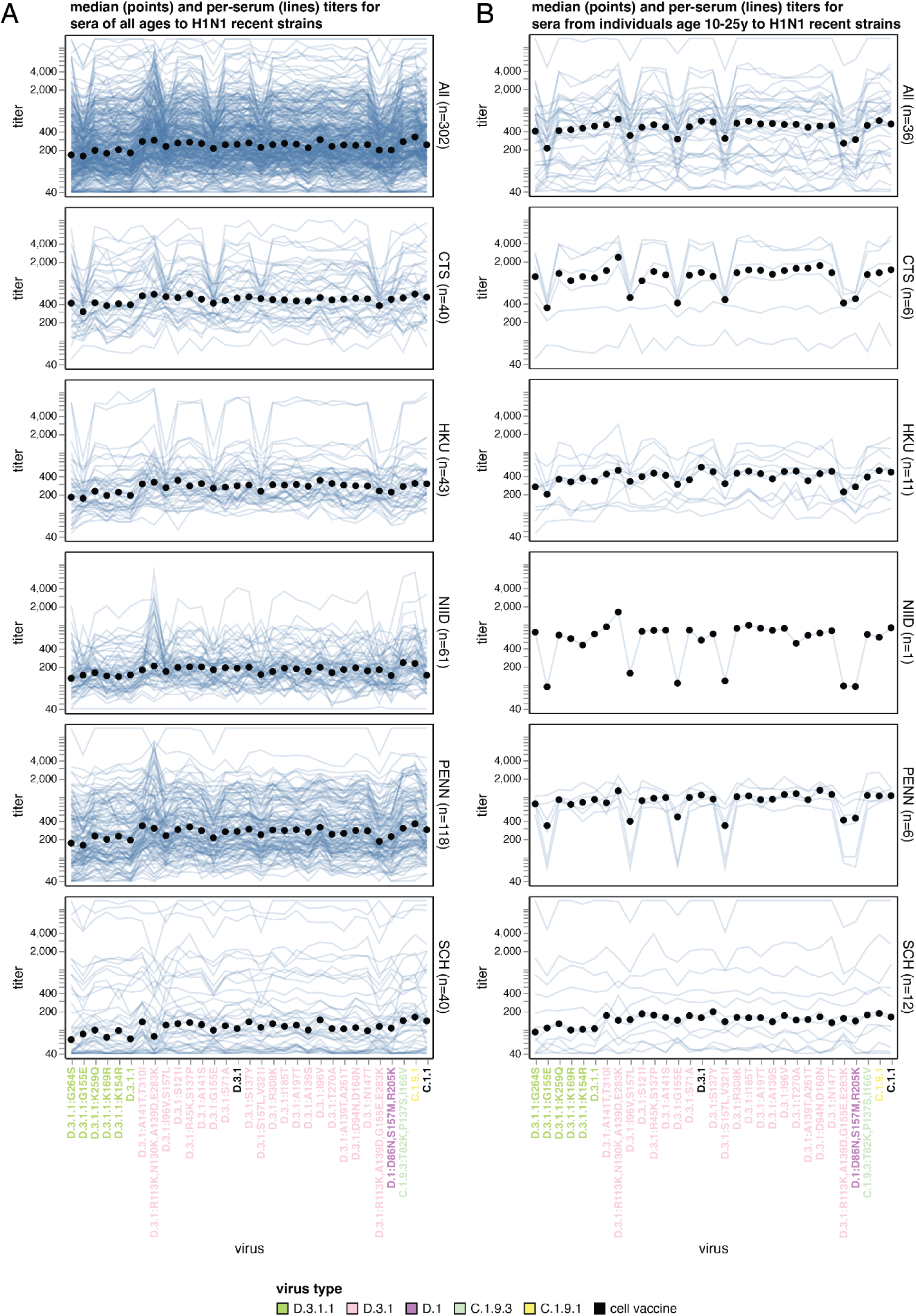
Human neutralizing antibody landscape against recent H1N1 viruses. Median titer across sera (black points) and titers for individual sera (blue lines) against the recently circulating H1N1 strains in the library for (A) sera from individuals of all ages and (B) sera from individuals between 10 to 25 years of age. In both panels, the top plot shows titers for all 302 sera, and the other plots show titers by serum set. The viral HA1 haplotype labels are colored by their subclade. The C.1.1 strain is the current cell-based 2025-2026 Northern Hemisphere vaccine strain, and a D.3.1 strain was chosen in September 2025 for the cell-based 2026 Southern Hemisphere vaccine (these strains are labeled in black). The dynamic range of our assays (determined by the dilution series used for the sera) enabled measurements of titers between 40 and 13619; sera with titers above or below this range are censored accordingly. See https://jbloomlab.github.io/flu-seqneut-2025to2026/human_H1N1_recent_individual_sera_vertical.html for an interactive version of this plot that can be subset on specific sera or age groups, and for which you can mouseover points and lines for details about viruses and sera. In particular, a slider at the bottom of this interactive plot allows you to subset on just sera from certain age ranges like is done in panel (B) of this figure. See https://jbloomlab.github.io/flu-seqneut-2025to2026/human_H1N1_recent_interquartile_range_vertical.html for a comparable plot showing the median and interquartile range. Note that this figure shows just the titers against the recent strains and the 2025-2026 and 2026 vaccine strains (31 strains total); see https://jbloomlab.github.io/flu-seqneut-2025to2026/ for plots that show titers against the older vaccine strains.

**Figure 7.**
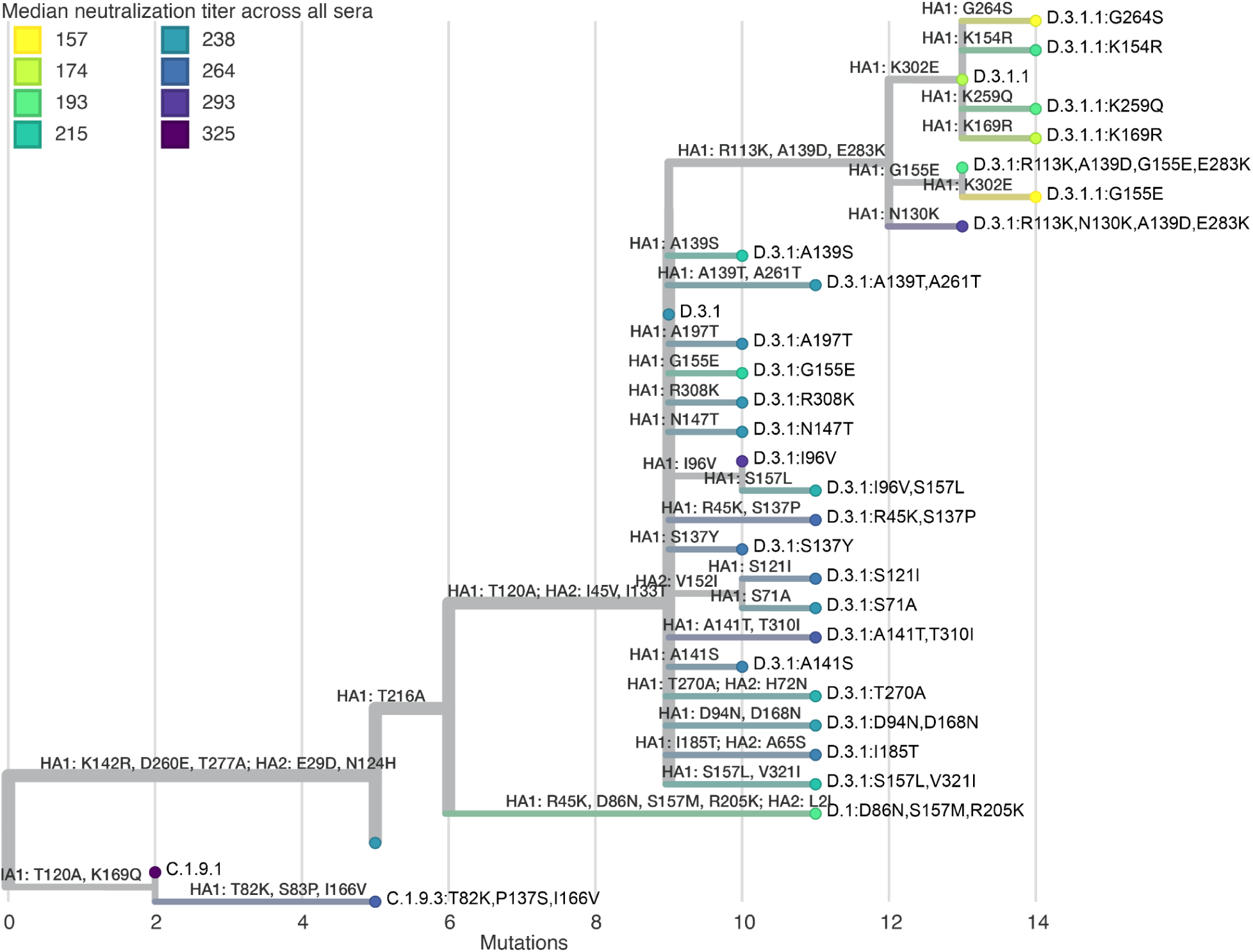
Phylogenetic tree of H1N1 HA proteins colored by median titer across all sera. Tree of H1N1 HA sequences in the library colored by the median titer across all sera. The tree is built on the protein sequences and branch lengths are amino-acid mutations relative to the root. All HA1 and HA2 mutations are labeled on branches. See https://nextstrain.org/community/jbloomlab/flu-seqneut-2025to2026@main/H1N1 for an interactive version of this tree with additional coloring options, a measurements panel with the individual serum titers, and an option to show all amino-acid mutations on branches.

There are striking strain-specific patterns in H1N1 strain neutralization for subsets of sera that are not well captured by the median titers across all sera (see **Figure 6A** and especially the interactive version linked in its legend). The most dramatic of these trends is that adolescents and young adults have clearly reduced titers to strains with mutations at sites 155 or 157 compared to individuals of other ages (**Figure 6B**). This observation parallels prior work that has found strong age variation in neutralization titers (and likely susceptibility to infection) of different age groups to H1N1 strains with specific HA mutations (Linderman et al. 2014; Petrie et al. 2016; Arevalo et al. 2020). Similarly, some sera from Seattle Children’s Hospital (SCH) have reduced titers to the D.3.1:R113K,N130K,A139D,E283K strain, whereas most adult sera show similar or higher titers to this strain as to other D.3.1 strains. This difference is likely attributable to site 130, which transitioned from K to N between 2018 to 2022, making N130K a novel mutation for children but a reversion to an amino-acid identity seen in previous exposures for adults.

### Changes in neutralization titers after vaccination with the Northern Hemisphere 2025 egg-based vaccine

Our measurements include pre- and 28-day-post-vaccination titers for 59 individuals following receipt of the 2025-2026 Northern Hemisphere Flulavel trivalent egg-based vaccine (the PENN sera set, see **Figure 3**). During the 2025-2026 Northern Hemisphere season, the egg-based vaccine components for H3N2 and H1N1 were a J.2 strain (A/Croatia/10136RV/2023) and a D strain (A/Victoria/4897/2022), respectively.

Across recent H3N2 strains, vaccination typically induced a moderate increase in titers after 28 days (median fold changes of ∼1.5 to 2.5), with some subclade-specific trends (**Figure 8A**). Both the final post-vaccination titers and the fold increase in titers tended to be lower for strains belonging to the most recent subclades (K, J.2.4 and J.2.3) (**Figure 8A**). There was appreciable variation in the fold-increase in titer across subclade K strains: the base K subclade had a ∼2-fold increase, which is on par with other clades, but other subclade K strains including those with mutations in antigenic region D (eg, at sites 96, 207, and 223) had lower vaccination-induced titer increases of only ∼1.5-fold. Note also that a few sera had much larger increases in titers post-vaccination than the median increase of ∼1.5 -to 2.5-fold, as can be seen from the interactive per-sera plots linked in the legend to **Figure 8**.

**Figure 8.**
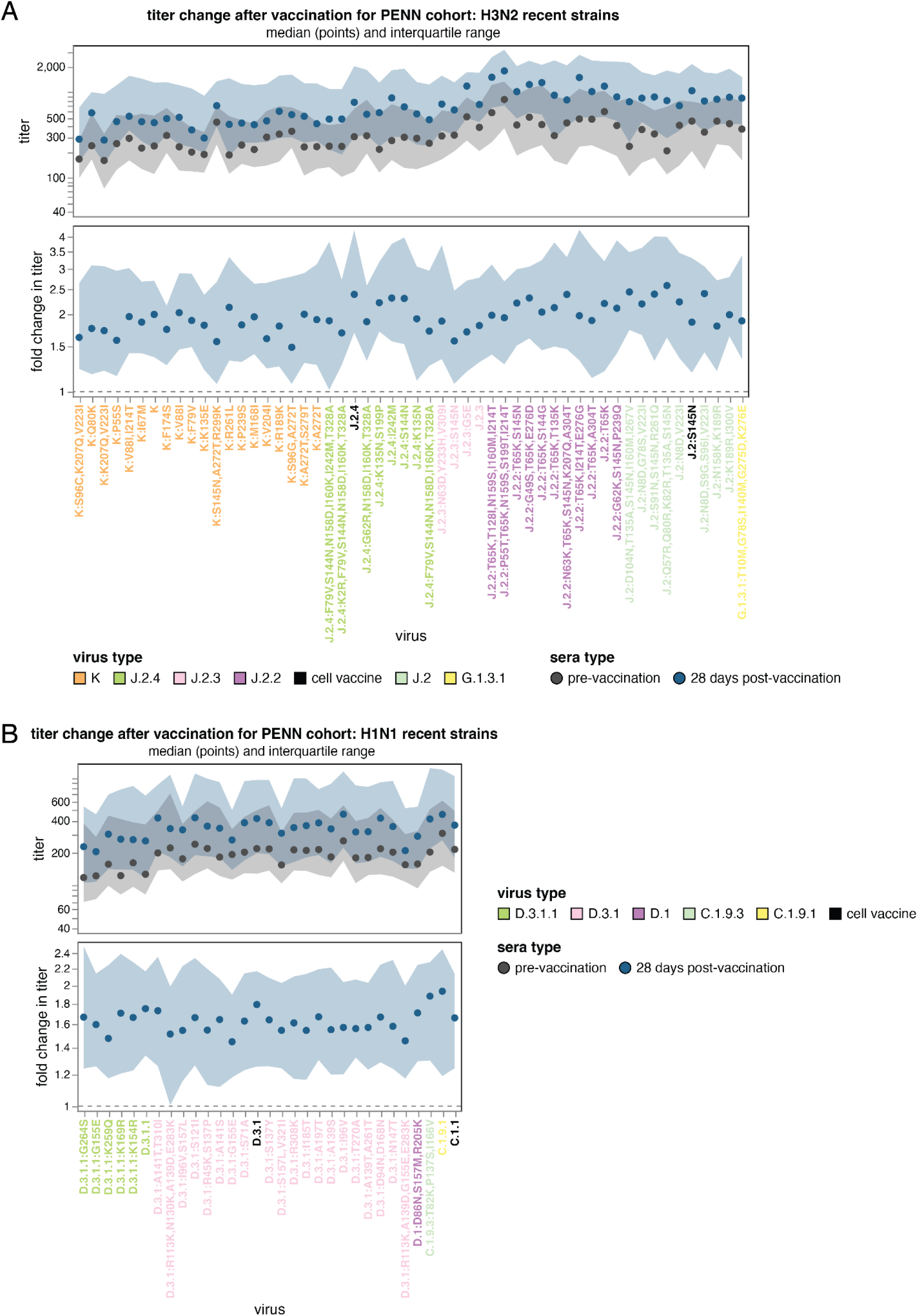
Change in human neutralizing antibody titers against recent H3N2 and and H1N1 strains after receipt of the 2025-2026 Northern Hemisphere influenza vaccine. The median titer (top) or fold-change in titer (bottom) across all sera in PENN set pre- and 28-day post vaccination (with a 2025-2026 egg-based vaccine) for all **(A)** H3N2 and **(B)** H1N1 strains in the library. The black points are the median across sera, and the shaded blue and black regions are the interquartile range. The viral haplotype labels are colored by their subclade or by status as a recent cell-based vaccine strain. See https://jbloomlab.github.io/flu-seqneut-2025to2026/PENN_pre_post_vax_H3N2_recent_interquartile_range_horizontal.html and https://jbloomlab.github.io/flu-seqneut-2025to2026/PENN_pre_post_vax_H1N1_recent_interquartile_range_horizontal.html for interactive versions of these plots. See https://jbloomlab.github.io/flu-seqneut-2025to2026/PENN_pre_post_vax_H3N2_recent_individual_sera_horizontal.html and https://jbloomlab.github.io/flu-seqneut-2025to2026/PENN_pre_post_vax_H1N1_recent_individual_sera_horizontal.html for comparable plots showing all individual sera.

For H1N1, the median increase in titers across sera are more modest (∼1.4-2-fold) than those observed for H3N2 (**Figure 8B)**. While pre-vaccination titers were lowest to the D.3.1.1 clades (the key clade that emerged in 2025-2026), post-vaccination titer increases to D.3.1.1 viruses are similar to those for strains in the D.3.1 subclade (**Figure 8B**). The weakest post-vaccination responses were observed in G155E-containing strains and in D.3.1.1:K259Q.

## Discussion

Here we have used high-throughput sequencing-based neutralization assays to measure the titers of a diverse set of human sera against a large set of HAs representing H3N2 and H1N1 influenza strains circulating as of early 2026. For both of these subtypes, new subclades (K for H3N2 and D.3.1.1 for H1N1) have become dominant in the six months since the 2026 Southern Hemisphere vaccine strains were selected. In concordance with other studies (Liu et al. 2026; Wang et al. 2026; Dee et al. 2026; Kirsebom et al. 2025; Separovic et al. 2026; Cheng et al. 2026; Ikonen et al. 2026), our measurements show that subclade K is generally more poorly neutralized by human serum antibodies than other strains, likely explaining their rise. Importantly, we also find that among subclade K H3N2 strains, those with mutations in antigenic regions D and E have especially low titers (Liu et al. 2026), raising the possibility that such strains could increase in frequency over the coming year.

In addition to these major trends, our data also shows additional fine-grain variation in titers to strains within the same subclades, and extensive heterogeneity in titers across sera, some of which partially stratifies with age group. How to best account for this additional heterogeneity in forecasting evolution and choosing vaccine strains remains an open question. By making this large dataset immediately available for analysis, we therefore hope both to inform vaccine-strain selection for the 2026-2027 season as well as spur further studies of how to leverage large and near-real-time neutralization data (Kikawa et al. 2025) to advance public health by improving understanding and forecasting influenza evolution.

Our study has several limitations. First, although we analyze a large set of sera, these sera are taken from individuals from just five different geographic locations and the vast majority of children sera were from Seattle Children’s Hospital. While major seasonal influenza strains generally circulate globally, it is possible that population-level influenza immunity may differ somewhat from region to region (Wen et al. 2016; Bedford et al. 2015). Additionally, most sera were collected before K subclade strains became dominant in late-November 2025, and so our measurements are more reflective of population influenza immunity at the start rather than the end of the 2025-2026 influenza season. Our experiments also only study strain-to-strain variation in neutralization of HA, but cellular immunity and antigenic variation in neuraminidase can also affect influenza evolution (Rosu et al. 2026; Sandbulte et al. 2011; Machkovech et al. 2015). Finally, our study does not analyze influenza B, which antigenically evolves slower than influenza A but still causes appreciable disease and is an important component of the annual influenza vaccine (Rosu et al. 2022; Akin et al. 2025).

Despite these caveats, the subclades of both H3N2 (subclade K) and H1N1 (subclade D.3.1.1) that our experiments measured to have the lowest neutralization titers spread to become dominant during the 2025-2026 Northern Hemisphere influenza season. This fact suggests that rapid large-scale measurements of neutralization titers provide substantial information relevant to forecasting viral evolution. Going forward, it will be important to assess if the more fine-grained data our study provided on which strains within subclades K and D.3.1.1 have the lowest neutralization titers prove useful for understanding which descendants of these subclades become dominant over the coming influenza season.

## Acknowledgements

We thank Zoe Munsey and Sophie Rotter-Aboyoun for assistance with collection of the SCH sera, and Eduard Grebe, Hasan Sulaeman, and Valerie Green for assistance with the collection of the CTS blood donor plasma. We also thank Catherine Jacob-Dolan, Sara Sunshine and Bernadeta Dadonaite for technical expertise and advice. This work was funded in part by the NIH/NIAID under R01AI165281 (to TB, JH, and JDB for the experimental work and data analysis performed at the Fred Hutchinson Cancer Center), 75N93021C00015 (to SEH, BJC, TB and JDB), F30AI186284 (to CK for the experimental work and data analysis performed at the Fred Hutchinson Cancer Center), R01AI165818 (to SAT and DJS), and 75N93021C00014 (to SAT and DJS). JDB and TB are Investigators of the Howard Hughes Medical Institute. This research was also supported by Dolores Covarrubias, Catherina Artikis, and the Genomics & Bioinformatics Shared Resource (RRID:SCR_022606) of the Fred Hutch/University of Washington Cancer Consortium (P30 CA015704) and by Fred Hutch Scientific Computing (NIH grants S10-OD-020069 and S10-OD-028685). SAT and DJS were funded in part by the UK Medical Research Council grant MR/Y004337/1 and Doctoral Training Grant. Sera from NIID were provided by Reiko Saito, with support from the Grants-in-Aid for Emerging and Reemerging Infectious Diseases from the Ministry of Health, Labour and Welfare, Japan (grant nos. 24HA2005). The EPI-HK study was also funded in part by the Theme-based Research Scheme under project no. T11-712/19-N (to BJC and NHLL) from the Research Grants Council from the University Grants Committee of Hong Kong. We gratefully acknowledge the authors and originating and submitting laboratories of the sequences from the GISAID EpiFlu Database(Shu and McCauley 2017) on which this research is partly based. This manuscript is the result of funding in part by the National Institutes of Health (NIH). It is subject to the NIH Public Access Policy. Through acceptance of this federal funding, NIH has been given a right to make this manuscript publicly available in PubMed Central upon the Official Date of Publication, as defined by NIH.

## Methods

### Data and code availability

All the analysis, code and documentation are publicly available on GitHub at https://github.com/jbloomlab/flu-seqneut-2025to2026. This repository is also archived on Zenodo at https://doi.org/10.5281/zenodo.20564730. That GitHub repository includes all details; key summary files are as follows:

- Information on the virus strains included in the library: **Supplementary File 1** and https://github.com/jbloomlab/flu-seqneut-2025to2026/blob/main/results/final_titer_data/human_viruses.csv
- Information on sera assayed against the library: **Supplementary File 2** and https://github.com/jbloomlab/flu-seqneut-2025to2026/blob/main/results/final_titer_data/human_sera.csv
- All neutralization titers after quality-control: **Supplementary 3** and https://github.com/jbloomlab/flu-seqneut-2025to2026/blob/main/results/final_titer_data/human_titers.csv
- Interactive summary plots showing median titers, individual serum titers, fraction of individuals below neutralization titer cutoffs, fold-change in titers post-vaccination are at the bottom sections of https://jbloomlab.github.io/flu-seqneut-2025to2026/. Earlier sections on that page also include details on quality control at the per-neutralization plate and per-serum level.
- Interactive Nextstrain proteins trees that can be colored by median titer or fraction of sera below titer cutoff (for all sera and for specific cohorts of sera) for H3N2 (https://nextstrain.org/community/jbloomlab/flu-seqneut-2025to2026@main/H3N2) and H1N1 (https://nextstrain.org/community/jbloomlab/flu-seqneut-2025to2026@main/H1N1)

### Biosafety

All experiments were performed at biosafety level 2. The work reported here involved influenza virions expressing HA ectodomain proteins from naturally-occurring recent human H3N2 or H1N1 influenza strains with non-HA genes from the lab-adapted A/WSN/1933 (H1N1) strain. Both recent human seasonal strains and the lab-adapted A/WSN/1933 strain are classified as biosafety level 2 by the CDC BMBL handbook (edition 6). This study did not involve any generation of viruses with non-natural HA amino-acid mutations; all HA ectodomains are identical to recent human seasonal viruses.

### Human sera and plasma

Serum and plasma samples were taken from children and adults across ages and geographical regions through a combination of epidemiological studies, vaccination cohorts, residual blood donation samples, and residual blood draws from hospitals.

The CTS plasma samples were obtained in November 2025 from blood donations collected across the USA from individuals 19-86 years of age through a biobank maintained at Creative Testing Solutions (CTS) by a collaboration between Vitalant Research Institute and the American Red Cross.

The HKU sera were obtained through the ‘<u>E</u>valuation <u>P</u>opulation Immunity in <u>H</u>ong <u>K</u>ong’ (EPI-HK) study, a community-based longitudinal observational cohort study of ∼2000 individuals run by the University of Hong Kong since 2020 (Cowling et al. 2022). For this study 48 sera were tested, including (1) 35 sera from the 35 out of 42 participants between 10 and 79 years of age who were tested in a similar study prior to the September 2025 vaccine-strain selection (Kikawa et al. 2025) and remained in the study, plus 7 sera from another 7 participants as replacement to those lost to follow up by matching on age (±10 years of birth year), sex, and vaccination history between 2020/21 to 2024/25 (winter) influenza seasons, with all 42 sera collected between July and November 2025; and (2) 6 sera from 4 participants with influenza virus infection identified during May to August 2025 from acute respiratory illness (ARI) active surveillance and confirmed by in-house PCR, including 4 sera from 2 participants with paired pre- and post-infection sera and 2 sera from 2 participants with pre-infection sera only. However, in the figures and results here we only report the titers for the 1 of these 6 post-infection sera collected no later than July 2025 (as we are focusing on relatively recent sera); the other titers from sera collected before July of 2025 are in the GitHub repository but not included in the results in this paper. The University of Hong Kong’s Institutional Review Board granted approval for the EPI-HK study protocol. All participants, or their legal guardians where applicable, provided written informed consent prior to enrollment.

The NIID sera were obtained from participants in a vaccine study by the National Institutes of Infectious Disease in Japan. These sera were from unique individuals 20-105 years of age at the day 0 pre-vaccination time point in November 2025.

The PENN plasma were from adults 24-81 years of age enrolled in a vaccination cohort study by the University of Pennsylvania on the day of and 28 days-post vaccination with the 2025-2026 Northern Hemisphere seasonal influenza vaccine (FluLaval trivalent influenza virus vaccine from GlaxoSmithKline) between October-November 2025. This study was approved by the University of Pennsylvania Institutional Review Board under protocol number 849398. Note that these sera were recently analyzed against a few H3N2 strains by hemagglutination-inhibition assays(Liu et al. 2026).

The SCH sera are deidentified pediatric sera obtained from children 0-15 years of age not known to be immunocompromised receiving routine medical care at Seattle Children’s Hospital in November 2025. These sera were obtained with a signed waiver of consent and the approval from the Seattle Children’s Hospital Institutional Review Board.

Before use in sequencing-based neutralization assays, all sera and plasma were treated with receptor-destroying enzyme II and heat-inactivated following a similar protocol as previously described (J. M. Lee et al. 2019b). Briefly, receptor-destroying enzyme II (Seikan) was resuspended in 20 mL PBS and passed through a 0.22 um filter. Next, 75 uL of resuspended and filtered receptor-destroying enzyme II was incubated with 25 uL of each sera or plasma (constituting 1:4 dilution) in 96-well PCR plates (BioRad) at 37°C for 2.5 hours and then 55°C for 30 minutes. Plasma samples were then transferred to 96-well V-bottom polypropylene plates (Corning) and spun at 2000g for 10 minutes before the cleared supernatant was transferred to new plates. All sera and plasma were used immediately or stored at -80°C until use.

### Choice of strains to include in the sequencing-based neutralization assays

Our goal in library design was to select HA strains representative of circulating H1N1 and H3N2 HA diversity in October-November 2025, and likely to remain representative of circulating diversity into the 2025-2026 Northern Hemisphere influenza season. We used Nextstrain(Hadfield et al. 2018) pdmH1N1 and H3N2 6-month builds available in October-November 2025 to identify all HA haplotypes, and selected strains from these lists of HA haplotypes. During this process, we aimed to choose ∼100 total strains that either had the relatively highest growth rates compared to other contemporaneously circulating strains (Abousamra et al. 2024) or contained HA mutations at either previously defined receptor-binding-site-adjacent and/or antigenic sites (Koel et al. 2013; Wolf et al. 2006; Caton et al. 1982) or HA mutations arising more than expected from the underlying mutation rate (Bloom and Neher 2023; S. A. Turner et al. 2026). This initial process selected 58 H3N2 and 48 H1N1 strains. The code for generating lists of HA haplotypes and metadata used to select strains is available at https://github.com/nextstrain/seasonal-flu/blob/02b5b69/notebooks/pick_library_strains.py. The documentation and code describing the design of the barcoded HA constructs is available at https://github.com/jbloomlab/flu-seqneut-2025to2026/tree/main/non-pipeline_analyses/library_design.

The library design also included the HAs from the component strains of both H3N2 and H1N1 seasonal vaccine strains. For H3N2, we included cell- and egg-based vaccine strains from the 2020 to the 2025-2026 Northern Hemisphere vaccine, and the cell-based vaccine for the 2026 Southern Hemisphere vaccine. For H1N1, we included cell- and egg-passaged vaccine strains from the 2018 to the 2025-2026 Northern Hemisphere vaccine, and the cell-based vaccine for the 2026 Southern Hemisphere vaccine. For H1N1 vaccine strains, most egg-passaged strains had to be dropped as they did not grow well in our system which grows viruses in mammalian cell lines.

Following library generation and quality control procedures, our final libraries comprised 186 barcodes spanning 53 recent H3N2 strains, 30 recent H1N1 strains, 4 historical H3N2 vaccine strains (egg-and cell-produced), and 4 historical H1N1 vaccine strains (egg- and cell-produced) (https://github.com/jbloomlab/flu-seqneut-2025to2026/blob/main/data/viral_libraries/flu-seqneut-2025to2026-barcode-to-strain-actual.csv). The initial library design included a slightly higher number of strains and barcodes, as some were excluded during library generation and quality control processes: 235 barcodes covering 58 recent H3N2 strains, 48 recent H1N1 strains, 4 historical H3N2 vaccine strains (egg-and cell-produced), and 4 historical H1N1 vaccine strains (egg- and cell-produced) (https://github.com/jbloomlab/flu-seqneut-2025to2026/blob/main/data/viral_libraries/flu-seqneut-2025to2026-barcode-to-strain-designed.csv).

### Cloning of barcoded HAs

The sequencing-based neutralization assays require inserting barcodes into the HA gene so that the assay can be read out by barcode sequencing (Kikawa et al. 2025; Loes et al. 2024; Kikawa et al. 2026). We used a previously described approach to insert the barcodes into the HA gene without disrupting viral genome packaging (Kikawa et al. 2025; Loes et al. 2024; Kikawa et al. 2026; Welsh et al. 2024). The 16-nucleotide barcodes sequences were randomly generated, but then checked to specifically avoid barcode sequences used in prior sequencing-based neutralization assays libraries (Kikawa et al. 2025; Loes et al. 2024; Kikawa et al. 2026) or that began with the nucleotides ‘GG’ (we have found such barcodes do not sequence well). All HA genes and linked barcodes were synthesized, cloned and sequence-verified by Twist Biosciences. As described previously (Kikawa et al. 2025, 2026; Welsh et al. 2024), the plasmid backbone for both the H1N1 and H3N2 constructs was the derivative of the pHH21 uni-directional reverse genetics plasmid (Neumann et al. 1999); see https://github.com/dms-vep/flu_h3_hk19_dms/blob/main/library_design/plasmid_maps/2851_pHH_WSNHAflank_GFP_H3-recipient_duppac-stop.gb for a map of this plasmid backbone. Exemplar plasmid maps for a H1 and H3 strain are at https://github.com/jbloomlab/flu-seqneut-2025/blob/main/non-pipeline_analyses/library_design/plasmids/example_constructs and the full set of all plasmid maps is at https://github.com/jbloomlab/flu-seqneut-2025to2026/tree/main/non-pipeline_analyses/library_design/construct_order/plasmids. The barcodes linked to each strain in the final library are at https://github.com/jbloomlab/flu-seqneut-2025to2026/blob/main/data/viral_libraries/flu-seqneut-2025to2026-barcode-to-strain-actual.csv.

### Generation and titration of barcoded viral libraries

To prepare the pooled libraries of barcoded virions used in the sequencing-based neutralization assays, each strain’s barcoded plasmids (note we had multiple replicate barcodes for most HAs) were pooled and then used to separately generate the barcoded virions with that particular HA. Exactly as described previously (Kikawa et al. 2025), the barcoded viruses expressing each different library HA were generated using reverse genetics (Neumann et al. 1999; Hoffmann et al. 2000) and then passaged on MDCK-SIAT1-TMPRSS2 cells. These virions contained the HA from the barcoded HA with a recent seasonal human influenza HA, and all the other genes from the lab-adapted A/WSN/1933 strain as described previously (Kikawa et al. 2025). Then, as in prior work (Kikawa et al. 2025), we made an equal-volume pool of the passaged viruses to determine the relative transcriptional titer of each virus. We used these relative transcriptional titers of each viral strain (see https://github.com/jbloomlab/flu-seqneut-2025to2026/blob/main/non-pipeline_analyses/library_pooling/notebooks/260107_equal_volume_pool.ipynb) to create a repooled library where each strains’ barcodes would be present roughly equally in the pool. As described previously, we repeated this experiment to affirm strains were roughly equally balanced, as well as determine the range of virus dilutions where viral transcription tracked linearly with the amount of virus particles added to cells (see https://github.com/jbloomlab/flu-seqneut-2025to2026/blob/main/non-pipeline_analyses/library_pooling/notebooks/260126_balanced_repool.ipynb). Based on this analysis, for the experiments described here we used a 1:24 dilution of the virus library infected on 1.5e5 MDCK-SIAT1 cells per well, as it was in the early part of this linear range where viral transcription is linearly correlated with viral neutralization.

### Sequencing-based neutralization assay

The protocol for the sequencing-based neutralization assays was identical to that outlined previously (Kikawa et al. 2025). A step-by-step protocol is available at https://dx.doi.org/10.17504/protocols.io.kqdg3xdmpg25/v2. Exactly as described previously, sera were diluted to 1:20 (accounting for the initial 1:4 dilution from the receptor-destroying enzyme treatment described above) in 50uL of influenza growth media (Opti-MEM supplemented with 0.1% heat-inactivated FBS, 0.3% bovine serum albumin, 100 ug/mL calcium chloride, 100 U/mL penicillin and 100 ug/mL streptomycin). This volume was then serially 2.3-fold diluted down 11 of the 12 columns of 96-well plate; the final column was used for the no-serum control wells required to normalize barcode counts to fraction infectivity (Loes et al. 2024). As determined in virus library titration experiments described above, virus library was then added to all wells at a 1:24 dilution in 50 uL, resulting in a range of serum dilutions from 1:40 to 1:13,619 across each 8-well column of a 96-well plate. These virus serum-mixtures were then incubated at 37°C with 5% CO2 for 1 hour before 1.5e5 MDCK-SIAT1 cells were added per well. After a 16 hour incubation, cells were lysed and barcodes were sequenced as described previously (Kikawa et al. 2025).

The sequencing data were analyzed as described previously using the *seqneut-pipeline* (https://github.com/jbloomlab/seqneut-pipeline), version 6.2.0. See the analysis configuration file (https://github.com/jbloomlab/flu-seqneut-2025to2026/blob/main/config.yml) for details about the parameters used for the barcode counting parameters, curve fitting and quality-control to remove low-quality barcodes, wells, and curves. For HAs with multiple barcodes, we report the median titer across barcodes. The computer code is available at https://github.com/jbloomlab/flu-seqneut-2025to2026/. This pipeline also generates HTML renderings of notebooks performing quality control and generating neutralization curves, as well as interactive plots summarizing the data, which can be explored at https://jbloomlab.github.io/flu-seqneut-2025to2026/.

### Phylogenetic analyses

The phylogenetic trees shown in **Figure 5** and **Figure 7** were built on HA protein sequences and then visualized via Nextstrain Community builds. The trees are rooted on older H3N2 and H1N1 sequences, and the branch lengths represent the number of amino-acid mutations. See https://github.com/jbloomlab/nextstrain-prot-titers-tree for the computer code used to build these trees; this computer code is embedded as submodule in the main GitHub repository for the project (https://github.com/jbloomlab/flu-seqneut-2025to2026).

Note that on the trees and in the figures, strains are labeled by their subclade designation (from a new dynamic nomenclature system (Neher et al. 2026)) plus any additional HA1 amino-acid mutations.

## Supplementary material

**Supplementary Figure 1.**
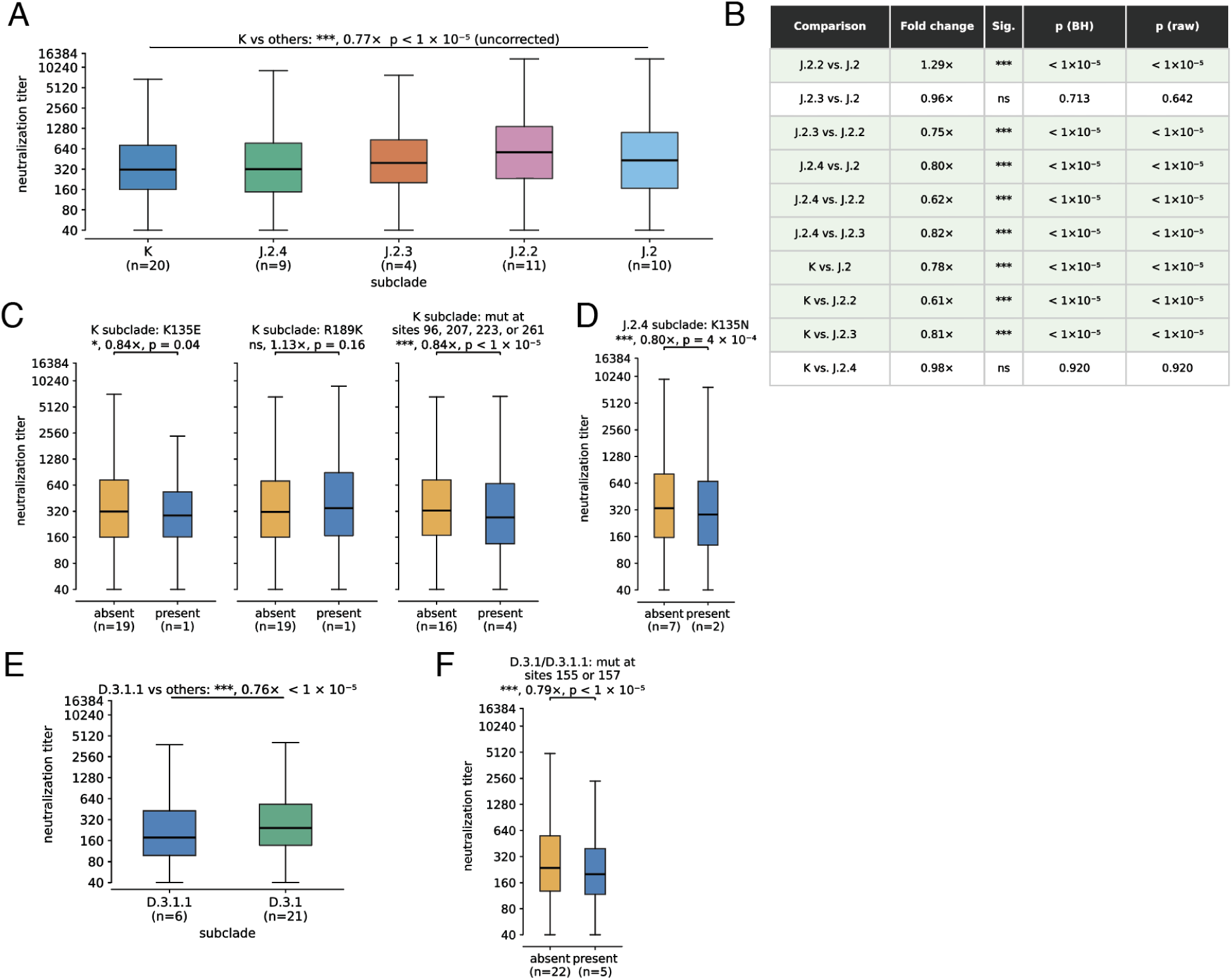
Statistical analysis on aggregated titers measured from 302 human serum samples. **(A)** Comparison of neutralization titers for all sera against all H3N2 subclade K viruses against all other 2025-2026-circulating strains in the library taken across all sera. The titers to K are significantly lower than the titers to other subclades (Mann-Whitney U test p < 1e-5). In the x-axis labels, n indicates the number of strains in that subclade in our library, and the box plots are the distribution of titers against those strains for all 302 sera. **(B)** All pair-wise comparisons between the different 2025-2026-circulating H3N2 subclades in the library with raw and Benjamini-Hochberg corrected p-values from the Mann-Whitney U test. Comparisons highlighted in green reached statistical significance. **(C)** Comparison of neutralization titers for all sera against all subclade K viruses with and without specific mutations or mutations at sets of sites in antigenic regions D and E, with Mann-Whitney p-values indicated. In the x-axis labels, n indicates the number of strains with that mutation. **(D)** The same as in **(C)**, but for subclade J.2.4 with and without K135N mutations. **(E)** The same as in **(B)**, but for H1N1 subclade D.3.1.1 against the other 2025-2026-circulating H1N1 subclade in the library. **(F)** The same as in **(C)** and **(D)**, but for H1N1 strains with and without mutations at sites 155/157.

**Supplementary File 1. CSV with details about the strains in the viral library**

Provides the strain name, collection date, subclade, derived haplotype, and nucleotide and protein HA ectodomain sequence for all viral strains in the library. Download or view at: https://github.com/jbloomlab/flu-seqneut-2025to2026/blob/main/results/final_titer_data/human_viruses.csv.

**Supplementary File 2. CSV with details about the tested sera**

Provides details about all sera assayed in the reported experiments. Download or view at: https://github.com/jbloomlab/flu-seqneut-2025to2026/blob/main/results/final_titer_data/human_sera.csv.

**Supplementary File 3. CSV with all neutralization titers reported here**

Provides neutralization titers measured by the reported experiments. Download or view at: https://github.com/jbloomlab/flu-seqneut-2025to2026/blob/main/results/final_titer_data/human_titers.csv.

